# Identification of *Escherichia coli* ClpAP in regulating susceptibility to type VI secretion system-mediated attack by *Agrobacterium tumefaciens*

**DOI:** 10.1101/734046

**Authors:** Hsiao-Han Lin (林筱涵), Manda Yu (余文廸), Manoj Kumar Sriramoju, Shang-Te Danny Hsu (徐尚德), Chi-Te Liu (劉啟德), Erh-Min Lai (賴爾珉)

## Abstract

Type VI secretion system (T6SS) is an effector delivery system used by gram-negative bacteria to kill other bacteria or eukaryotic host to gain fitness. In *Agrobacterium tumefaciens*, T6SS has been shown to kill other bacteria such as *Escherichia coli*. Interestingly, the *A. tumefaciens* T6SS killing efficiency differs when using different *E. coli* strains as recipient cells. Thus, we hypothesize that a successful T6SS killing not only relies on attacker T6SS activity but also depends on recipient factors. A high-throughput interbacterial competition assay was employed to test the hypothesis by screening for mutants with reduced killing outcomes caused by *A. tumefaciens* strain C58. From the 3909 *E. coli* Keio mutants screened, 16 candidate mutants were filtered out. One strain, Δ*clpP*::Kan, showed ten times more resistant to T6SS-mediating killing but restored its susceptibility when complemented with *clpP in trans*. ClpP is a universal and highly conserved protease that exists in both prokaryotes and eukaryotic organelles. In *E. coli*, ClpP uses either ClpA or ClpX as an adaptor for substrate specificity. Therefore, the susceptibility of the Δ*clpA*::Kan and Δ*clpX*::Kan was also tested. The T6SS attack susceptibility of Δ*clpA*::Kan is at the same level as Δ*clpP*::Kan, while Δ*clpX*::Kan showed no difference as compared to that of wild-type *E. coli* BW25113. The data also suggest that ClpA-ClpP interaction, rather than its protease activity, is responsible for enhancing susceptibility to T6SS killing. This study highlights the importance of recipient factors in determining the outcome of T6SS killing.

---

In all ecosystems, interactions of the microbial communities have ramifications in affecting its macroscopic world. For example, gut microbiota can influence human health, while rhizosphere microbiome has a significant impact on plant health (1,2). Microbial interactions studies provide new insights to both the microenvironment and macroscopic world. Among various kinds of microbial interactions, competition is the dominant force in evolution (3,4). Microbial competition can lead to reducing access to nutrients and space of their competitors, disruption of aggressive phenotypes, or competitor elimination (4). The type VI secretion system (T6SS) is a common strategy that gram-negative bacteria use to eliminate its competitors; more than a quarter of the sequenced gram-negative bacteria harbors T6SS gene cluster homologs in its genome (5).

The mechanism of T6SS to eliminate its competitors is to inject toxic effector proteins into the competitor cells. Effector-producing cells also harbor its cognate immunity protein to prevent self-intoxication. The effector-immunity (E-I) genes are usually, if not always, found in pairs in the genome (6). The process of T6SS-dependent competitor elimination is called T6SS antibacterial activity or T6SS killing. In addition, the T6SS effectors, in some cases, can toxify its eukaryotic hosts, making T6SS a weapon with a broad range (7). The ubiquitous existence and the broad recipient range characteristics make T6SS a potential tool to fight against pathogens. For example, the number of plant pathogen *Xanthomonas campestris* was lower when co-cultured with biocontrol agent *Pseudomonas putida* only when *P. putida* harbors a functional T6SS *in planta* (8). It has also been suggested that supplying probiotic commensal bacteria with T6SS may have biomedical applications (9). Therefore, T6SS has been viewed as one of the promising strategies not only to fight against multidrug-resistant pathogens but also could also serve as an eco-friendly biopesticide.

To maximize the effect of pathogen killing, comprehensive understanding of T6SS would be essential. The magnitude of T6SS killing is often evaluated by interbacterial competition assay in a contact-dependent manner (10,11). The interbacterial competition outcome is usually determined by two factors, the competition environment and the competitor (10,12). For the competition environment, it has been shown that T6SS killing is more pronounced in the environment which is similar to or mimic their ecological niches. For example, T6SS antibacterial activity of plant pathogen *Agrobacterium tumefaciens* against *Pseudomonas aeruginosa* was only observed *in planta* but not *in vitro* on an agar plate (12). In entero-pathogen *Salmonella enterica* serovar Typhimurium, T6SS killing against *E. coli* was enhanced when the environment contains bile salt (13). While there is a wealth of knowledge in regulation and action of T6SS activity of attacker cells, much less is understood about how and what recipient cell factors affect the outcome of T6SS killing.

Bacteria can utilize its T6SS to kill one type of bacteria but not another. For examples, *A. tumefaciens* was able to antagonize *E. coli* but not its susceptible siblings when co-cultured at an acidic condition where T6SS secretion is active (12,14). In contrast, such intra-species antagonism can occur *in planta* (14,15). These results suggest that the failure of the intra-species antagonism was not due to insusceptible siblings and that the recipient genetic features might also play a significant role in T6SS killing outcome. It was proposed that *Pseudomonas aeruginosa* may be able to hijack the elongation factor thermo-unstable (EF-Tu) protein of the recipient cell to grant access of its Tse6 effector into the recipient cytoplasm (16). However, another study demonstrated that Tse6 could penetrate the double bilayer of the EF-Tu-free liposome and exerted its toxicity inside it (17), making the role of EF-Tu involved in interbacterial competition of Tse6 elusive. Recently, a T6SS study in *Serratia marcescens* demonstrated that the recipient protein DsbA plays a role in activating *S. marcescens* T6SS effectors Ssp2 and Ssp4 but not Rhs2 (18). The *S. marcescens* T6SS kills its Ssp2-sensitive siblings only when the recipient cells harbor *dsbA* homologs (*dsbA1*^+^ *dsbA2*^+^). The same results were also observed using Ssp4-sensitive but not Rhs2-sensitive strain as a recipient cell. The results highlight the necessity of recipient DsbA to activate specific T6SS effectors. As DsbA is not an effector nor an immunity protein, this study also demonstrates that recipient genetic features other than E-I pairs affect the outcome of T6SS killing. However, the recipient genetic factors other than E-I pairs affecting the outcome of T6SS antibacterial activity have not been studied thoroughly.

This study aimed to explore the recipient genetic factors that affect the T6SS killing outcome using the well-characterized T6SS-possessing plant pathogen *A. tumefaciens*, a causative agent of crown gall disease in many different plants. The *A. tumefaciens* C58 harbors three effector proteins: type VI DNase effector 1 (Tde1), Tde2, and type VI amidase effector (Tae). The Tde proteins are the main contributor to *A. tumefaciens* T6SS-dependent interbacterial competition (12). We report here a high-throughput, population level, interbacterial competition screening platform for identifying the recipient genetic factors that are contributing to *A. tumefaciens* C58 T6SS killing outcome. We identified 16 candidates using this system, and four of them were confirmed to play a role by complementation *in trans*. One of the confirmed genes, *caseinolytic protease P* (*clpP*), was highlighted in this study due to the prominent phenotype. A functional ClpP complex consists of a tetradodecameric ClpP and its associated AAA^+^ ATPase substrate recognizing partner ClpA or ClpX (19). Further mutant studies show that *clpA*, but not *clpX*, is involved in the outcome of *A. tumefaciens* T6SS killing. Interestingly, our data suggest that the ClpAP complex formation, rather than ClpP protease activity, affects the outcome of T6SS killing. This work not only provides a new screening platform for elucidating factors for competitor eliminating systems but also strengthens the importance of recipient genetic factors in the outcome of T6SS antibacterial activity.

## Results

### The T6SS-dependent killing outcome differs between E. coli DH10B and BW25113

While searching for the recipient genetic factors that influence the T6SS killing outcome of *A. tumefaciens*, we optimized the competition medium (named as *<u>A</u>grobacterium* <u>K</u>ill-triggering, AK medium) for enhancing killing activity that characterized by only providing minimal minerals at pH 5.5. The AK medium contains basic minerals at pH 5.5 and provides an ideal condition to screen the *E. coli* mutants with enhanced resistance to *A. tumefaciens* T6SS killing. We first observed that the *A. tumefaciens* C58 T6SS killing to two different *E. coli* strains, BW25113 and DH10B, on AK medium. The recovered growth of strain BW25113 was always lower than that of strain DH10B when co-cultured with wild type *A. tumefaciens* C58 (hereafter referred to as wt) (Fig. 1A). Meanwhile, the survival of the two *E. coli* strains was not discriminable when co-cultured with Δ*tssL A. tumefaciens* C58 (hereafter referred to Δ*tssL*), a T6SS secretion-deficient mutant (Fig. 1A).

**Figure 1.**
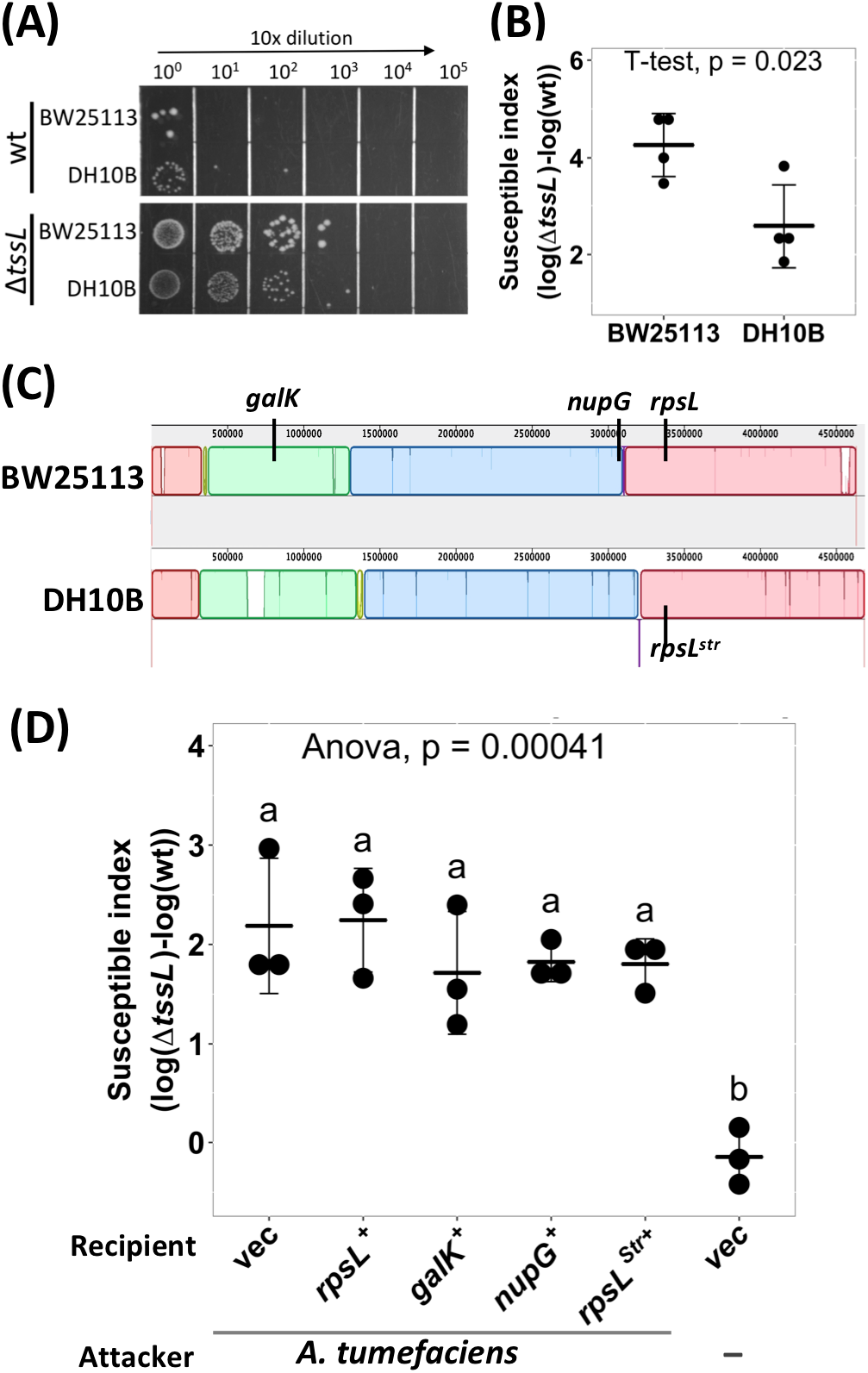
*A. tumefaciens* T6SS-dependent antibacterial activity against *E. coli* strains. **(A) and (B)** *A. tumefaciens* T6SS antibacterial activity against *E. coli* strains DH10B and BW25113. *A. tumefaciens* was co-cultured at a ratio of 30:1 with *E. coli* DH10B or BW25113, both *E. coli* strains harboring vector pRL662, on AK agar medium for 16 h. The bacterial mixtures were serially-diluted and spotted **(A)** or quantified by counting cfu **(B)** on gentamicin-containing LB agar plates to selectively recover *E. coli*. **(C)** Whole genome comparison between the chromosome of *E. coli* strain BW25113 (GenBank Accession Number CP009273) and DH10B (GenBank Accession Number CP000948) using Mauve (40). The BW25113 genome set as a reference. Colored block outlines the regions with homology to the reference without sequence rearrangement. The darker vertical lines within the box represent the region with less homology. Areas without color are regions that could not be aligned. **(D)** *E. coli* DH10B was complemented by either vector only (vec) or derivative expressing *rpsL*, *galK*, *nupG*, or *rpsL^Str^ in trans* before subjected to *A. tumefaciens* T6SS-dependent antibacterial activity assay as described in (B). Susceptible Index (SI) was defined as the subtraction difference of the recovery log(cfu) of that attacked by Δ*tssL* to that attacked by wild type *A. tumefaciens* C58. Data are mean ± SD of three independent experiments calculated by t-test with *P* < 0.05 for statistical significance **(B)** or single factor analysis of variance (ANOVA) and TukeyHSD, in which two groups with significant differences are indicated with different letters (a and b) **(D)**.

For more intuitive readout, T6SS-dependent susceptible index (SI) was introduced to quantify the T6SS killing outcome. The SI was designated as the logarithm recovered colony-forming unit (cfu) of that attacked by Δ*tssL* subtracted by that attacked by wild type *A. tumefaciens*. The high SI value indicates strong *A. tumefaciens* T6SS killing. Consistent with serial dilution results, the mean SI between *A. tumefaciens* and BW25113 was significantly higher than that between *A. tumefaciens* and DH10B with *p*-value (T ≤ t, two-tail) of 0.02 (Fig. 1B), suggesting that some genetic factors of BW25113 may enhance *A. tumefaciens* C58 killing outcome in a T6SS-dependent manner. We tested whether the genes that are functional in BW25113 but not in DH10B could be the candidates. The *galK* and *nupG* are functional in BW25113 but are pseudogenes in DH10B, and the *rpsL* has a mutation in DH10B (*rpsL^Str^*), which render the strain resistant to streptomycin, but not in BW25113 (Fig. 1C). Thus, *rpsL*, *galK*, or *nupG* from BW25113 was cloned into pRL662 and expressed by constitutive *lacZ* promoter in DH10B as recipient for T6SS interbacterial competition assay (Fig. 1D). The DH10B expressed with either the *rpsL^Str^* or empty vector served as the negative controls. A group without attacker was also included to monitor whether the decrease in cfu after the competition solely comes from co-culture with *A. tumefaciens* attacker. The SI was not significantly different between DH10B and any of the complemented groups, and each had SI mean of about 2 (Fig. 1D). As there are still many different genes between BW25113 and DH10B, according to whole-genome comparison (Fig. 1C), it was not practical to test them one by one. The above results also imply that the genetic factors of BW25113 other than those mutated in DH10B may play a role in enhancing *A. tumefaciens* C58 T6SS killing.

### Establish a high-throughput population-level, interbacterial competition screening platform to identify E. coli mutants with less susceptibility to A. tumefaciens C58 T6SS killing

As the genetic factors of BW25113 that play a role in enhancing *A. tumefaciens* C58 T6SS killing cannot be readily identified, we screened the BW25113 single-gene mutant library (Keio collection from NBRP (NIG, Japan): *E.coli*) for strains with less susceptibility to *A. tumefaciens* T6SS-mediated killing. A regular interbacterial competition assay starts from culture and mixing the attacker cells and the recipient cells (Fig. 2A). Competition outcome is scored by spotting cocultured bacterial cells with 10 times serial dilution and by counting recovered cfu of recipient *E. coli* on selective media.

**Figure 2.**
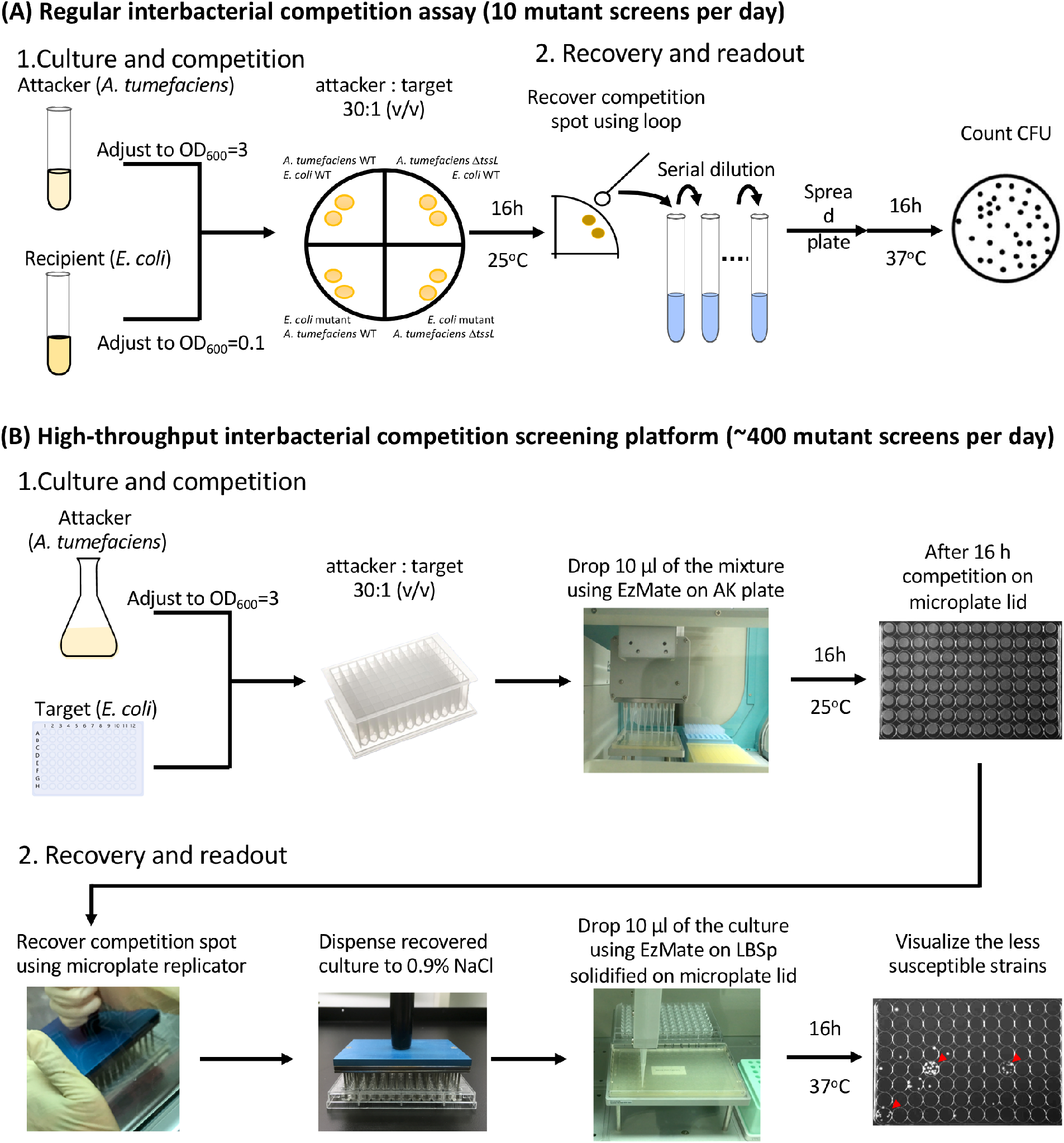
The high-throughput interbacterial competition screening platform. **(A)** Regular interbacterial competition assay. Cultured attacker *A. tumefaciens* and recipient *E. coli* were mixed and then spotted on the AK agar medium to allow interbacterial competition for 16 h at 25°C followed by recovery of mixed cultures, serial diluted, and then spread onto LB plate supplemented with appropriate antibiotics to select for recipient cells. **(B)** High-throughput interbacterial competition screening platform. Recipient cells were grown and mixed with attacker *A. tumefaciens* in a 96-well plate. The bacterial mixture was dropped onto the AK agar medium competition surface using an automated pipetting system. The competition surface was made on a microplate lid. Recovery was performed using microplate replicator. The candidates are the strains that show multiple colonies grown after recovery as opposed to wild type controls and most strains with no or few colonies. This high-throughput *A. tumefaciens* T6SS killing platform enables ~400 mutant screens per day.

In practice, a regular interbacterial competition assay (Fig. 2A) enables screening of ten mutants per day; which is not applicable as the Keio library contains 3909 strains. Therefore, we developed a high-throughput, population-level interbacterial competition screening platform that enables 96 simultaneous mutant screening (Fig. 2B). The recipient Keio *E. coli* strains were cultured in the 96-well, and the attacker *A. tumefaciens* was cultured in a flask. After the culture, the attacker *A. tumefaciens* was adjusted to OD_600_ equals to 3.0, then dispensed to a 2.2 mL deep-well plate. The recipient cells were added into the attacker-containing plates in a volume ratio of 30 to 1. Ten microliters of the attacker-recipient mixture were then dropped on the solidified competition agar on the 96-well lid with the automatic pipetting system. The cells were recovered after 16 h incubation at 25 °C then the cells were recovered. Microplate replicator was used to stamp onto the competition surface followed by recovery in the saline buffer (0.9% NaCl) to make the recovery area of each group consistent. The recovered bacteria were then spotted on kanamycin-containing LB agar solidified on microplate lid. The above competition condition enables *A. tumefaciens* to effectively kill wild-type BW25113 recipient so that only a few or no cells would survive, which made the readout of the mutant candidates simple—the ones with the multiple colonies are the candidates (Fig. 2B).

All the 3909 strains in the Keio were screened using C58 as the attacker. In each screening, at least two wild type *E. coli* BW25113 were incorporated and screened in parallel as parental control. The Keio mutants that formed colonies in this stage were selected, and 196 strains were identified. To further confirm their phenotype, the 196 candidates were subjected to second screening using both C58 and Δ*tssL* as the attacker. Sixteen Keio mutants were ten times less susceptible to C58 killing and showed no significant difference to that of wild type *E. coli* when co-cultured with Δ*tssL*.

### Confirmation of the E. coli mutants with less susceptibility to A. tumefaciens C58 T6SS killing

After the screening, 6 out of the 16 candidates were selected and verified by both regular interbacterial competition assay (Fig. 2A) and by complementation test (Table 1, Fig. S1). For complementation, wild type gene from BW25113 was cloned into plasmid pTrc200HA plasmid and expressed by *trc* promoter by IPTG induction in respective mutants for T6SS interbacterial competition assay. Representative results of the regular interbacterial competition assay using one of the candidates, Δ*clpP*::Kan (labeled as Δ*clpP*), and its complemented strain *clpP*^+^ are shown in Fig. 3A. After 16 h competition using wild type *A. tumefaciens*, the recovered cfu of Δ*clpP* was about 10^4^ while it was about 5×10^2^ in both wild type BW25113 or *clpP*^+^ (Fig. 3A). The initial cfu of the co-cultured *E. coli* recipient cells at 0 h was about 10^6^ in all groups (one-way ANOVA with *P* = 0.98, Fig. 3A), indicating that the *E. coli* cfu difference in wt co-cultured group was not due to different initial bacteria titer. When using *A. tumefaciens* Δ*tssL* as the attacker, the cfu at 0 h, 16 h, and as well as among different recipient *E. coli* strains was not significantly different (Fig. 3A). This result indicates that the Δ*tssL* had no competitor eliminating ability. If expressed as susceptibility index, the mean SI of the BW25113 wild type to *A. tumefaciens* C58 is 3.2, which is 1.8-fold higher than that of Δ*clpP* (SI=1.7) (Fig. 3B). The less susceptible phenotype of the Δ*clpP* can be fully complemented *in trans* (*clpP*^+^) (Fig. 3). These results confirmed that *clpP* contributes to enhancing susceptibility to T6SS antibacterial activity of *A. tumefaciens* C58.

**Figure 3.**
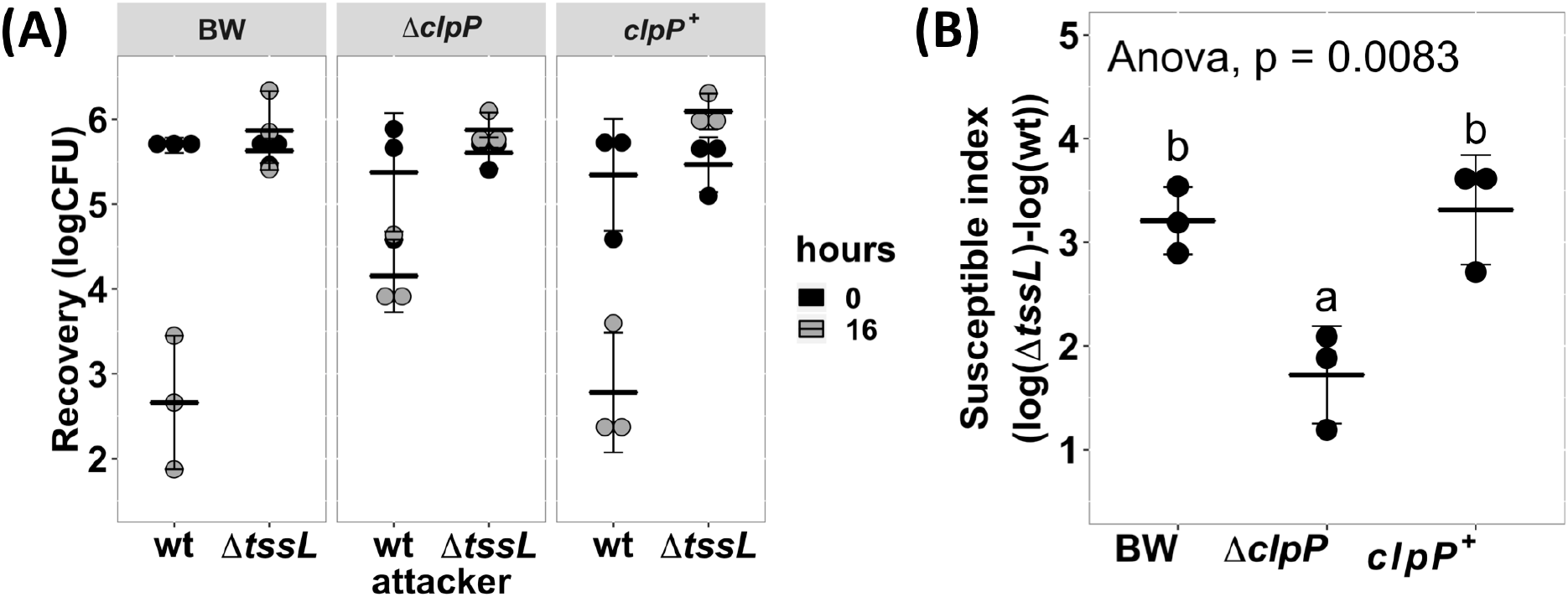
*A. tumefaciens* susceptibility to T6SS-dependent antibacterial activity was reduced in *E. coli clpP::kan* and can be fully complemented *in trans*. **(A)** Recovery of surviving *E. coli* cells at 0 h and 16 h after co-cultured with either *A. tumefaciens* wild type C58 (wt) or Δ*tssL* (Δ*L*) at a ratio of 30:1. **(B)** The susceptible index (SI) of *E. coli* BW25113 wild type (BW), Δ*clpP*, and Δ*clpP* complemented with *clpP* expressed on plasmid (*clpP*+) was calculated from the recovery rate shown in (A). Statistical analysis involved single factor analysis of variance (ANOVA) and TukeyHSD. Data are mean ± SD of three independent experiments and two groups with significant differences are indicated with different letters (a and b) (*P* < 0.05 for statistical significance).

**Table 1.** *E. coli* strains that showed reduced susceptibility to *A. tumefaciens* T6SS attack.

| No. | Resource (JW ID) | disrupted gene | Gene products affected by Kanamycin cassette insertion<sup>a</sup> | reduced susceptibility<sup>b</sup> | trans complementation<sup>c</sup> |
| --- | --- | --- | --- | --- | --- |
| 1 | JS0427 | *clpP* | ClpAXP, ClpXP, ClpAP | O | O |
| 2 | JW0710 | *gltA* | citrate synthase | O | O |
| 3 | JW1658 | *ydhS* | FAD/NAD(P) binding domain-containing protein YdhS | O | O |
| 4 | JW1346 | *ydaE* | Rac prophage; zinc-binding protein | O | Δ |
| 5 | JW0985 | *cbpA* | curved DNA-binding protein | O | X |
| 6 | JW1792 | *yeaX* | carnitine monooxygenase | X | n.d. |

<sup>a</sup> Gene products information was obtained from the EcoCyc database (41)
<sup>b</sup> Mutant strains with reduced susceptible index (SI) and showed significant difference under $P < 0.05$ was labelled as O, with no significant difference to that of wild type was labelled in X.
<sup>c</sup> Plasmid-born gene that can fully complement the disrupted gene is labelled in O, partially complemented is labelled in Δ, and cannot be complemented labelled in X. n.d.: not determined.

Of the six verified candidate mutant strains, five of them showed significant lower SI (*P* < 0.05) than that of wild type. These strains are *clpP*, *gltA*, *ydhS*, *ydaE*, and *cbpA*, the *cbpA* showed a milder reduction. One of the five strains, Δ*cbpA*::Kan, that show significant lower SI and could not be complemented *in trans* under the condition tested. As *cbpA* is the first gene in its operon, the significantly reduced SI could result from the polar effect of the mutant. Alternatively, the failure to complement may result from inappropriate expression level under the IPTG induction condition used during the interbacterial competition assays. One of the candidates, the Δ*yeaX*::Kan, did not differ in SI when compared to that of the wild type after conventional antibacterial assay verification (Table 1). Overall, the results demonstrated that the high-throughput, populational-level interbacterial competition screening platform is useful in identifying the recipient genetic factors that participate in T6SS killing but also verify and confirmed the recipient genetic factors that are important in *A. tumefaciens* T6SS killing.

### The ClpP protein but not its protease activity plays a critical role in enhancing susceptibility to A. tumefaciens T6SS killing

The Δ*clpP* is one of the strains that is less susceptible to *A. tumefaciens* T6SS-mediated antibacterial activity (Table 1, and Fig. 3). The ClpP complex is a well-studied, house-keeping AAA^+^ serine protease in *E. coli* (20–22). A functional ClpP complex consists of a tetradodecameric ClpP and its associated AAA^+^ ATPase substrate recognizing partner ClpA or ClpX, both in a hexameric form (19). The protease catalytic triad of the *E. coli* ClpP (*Ec*ClpP) is composed of S111, H136, and D185 (counted from the Met1) (23,24). Therefore, we asked whether the recipient cell ClpP protease is essential in enhancing *E. coli* susceptibility to *A. tumefaciens* C58 T6SS attack by using *E. coli* recipient strains Δ*clpP* complemented with pTrc200HA expressing either wild type or catalytic mutated ClpP S111A, H136A, and D185A. All ClpP variants contain a C-terminal HA tag. Surprisingly, two of the catalytic mutants S111A^+^ and H136A^+^ failed to complement, whereas the third catalytic mutant D185A^+^ can fully complement the phenotype (Fig. 4A).

**Figure 4.**
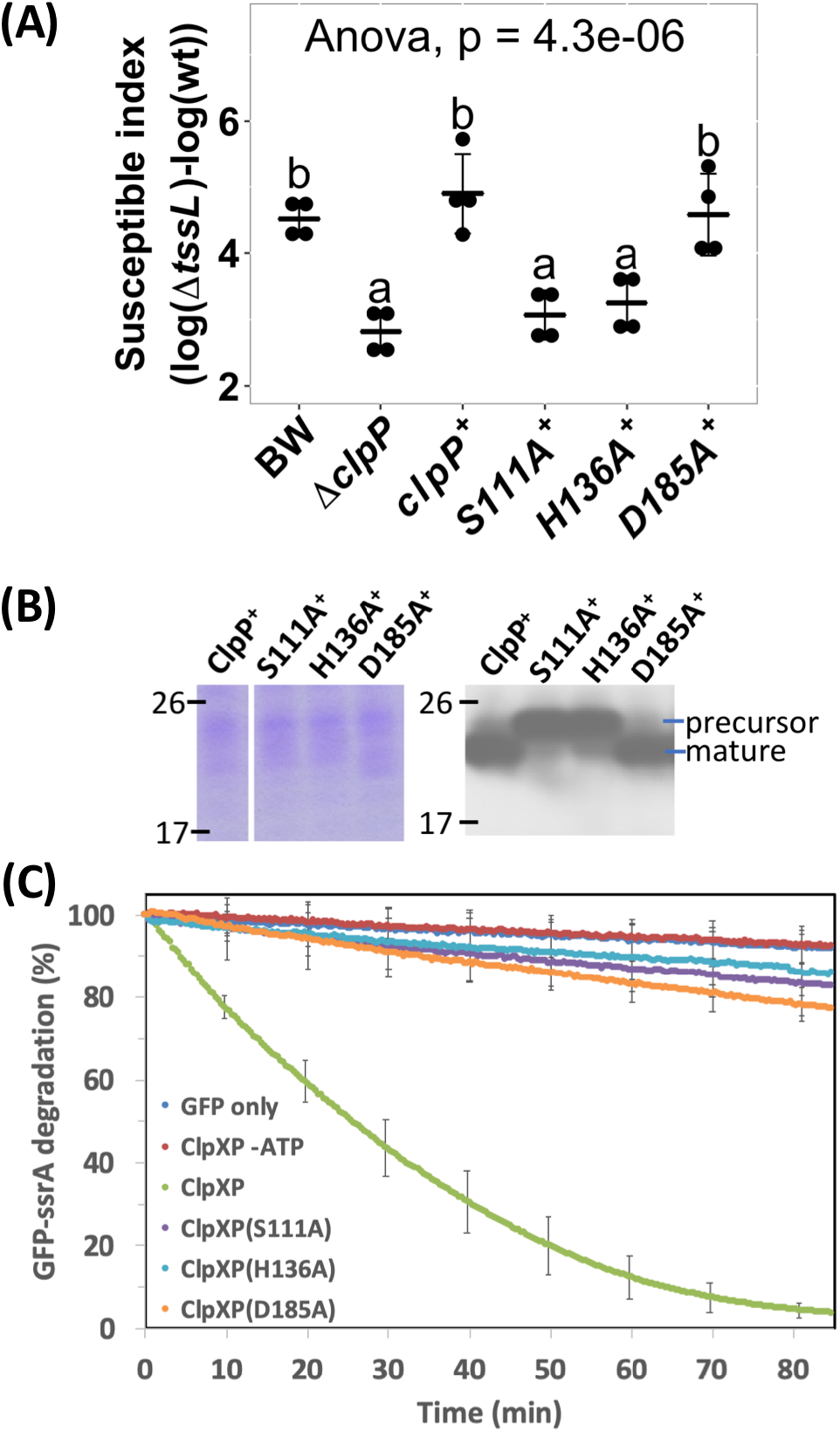
ClpP protease activity may not play an important role in enhancing *A. tumefaciens* T6SS antibacterial activity. **(A)** The susceptible index calculated from *A. tumefaciens* interbacterial activity assay against *E. coli*. The *A. tumefaciens* C58 wild-type or Δ*tssL* were co-cultured at a ratio of 10:1 with *E. coli* BW25113 wild type (BW), Δ*clpP*, and Δ*clpP* complemented with wild type *clpP* expressed on plasmid. The complemented *clpP* were either wild type or catalytic mutants ClpP_S111A_, ClpP_H136A_, and ClpP_D185A_ with C-terminus HA-tag. The susceptible index (SI) of each *E. coli* was calculated from the logarithm recovery rate of the Δ*tssL* co-cultured group minus that of the wild-type co-cultured group. Data are mean ± SD of four biological replicates from two independent experiments. Statistical analysis involved single factor analysis of variance (ANOVA) and TukeyHSD with *P* < 0.05 for statistical significance. **(B)** The ClpP protein levels of the Δ*clpP* complemented strains used in (A). The ClpP-expressing *E. coli* strains were cultured at the same condition used in interbacterial competition assay. Instead of co-culture with *A. tumefaciens*, protein samples were collected, normalized, and subjected to Western blot analysis of ClpP::HA and its variants. Three independent experiments were performed with similar results. **(C)** Protease activity assay of the ClpP and its catalytic mutant variants. The ClpP and its catalytic mutated variants was pre-assembled with ClpX followed by providing its substrate, the ssrA-tagged GFP. The GFP fluorescent signals were monitored over time. Data are mean ± SD of three biological replicates from one representative result, and at least two independent experiments were performed with similar results.

The difference of ClpP catalytic mutants to complement Δ*clpP* is not due to their protein-expression level as determined by Western blot, using anti-HA to detect the C-terminal HA-tag of the complemented ClpP, showed similar protein levels among the three catalytic mutants (Fig. 4B). However, the protein bands of the ClpP_S111A_ and ClpP_H136A_ migrated significantly slower than that of the ClpP_wt_ and ClpP_D185A_, which is consistent with previous findings (24,25). A functional ClpP is processed to form a mature form with the removal of the N-terminus 1-14 amino acid. Both ClpP_S111A_ and ClpP_H136A_ protein is defective in processing into the mature form and therefore accumulates as fulllength propeptide form while ClpP_D185A_ is processed into mature form (24,25).

To ensure the catalytic site mutants are indeed deficient in protease activity, we performed the ClpP protease activity of these mutants. We took advantage of the widely adopted ClpP protease degradation assay using GFP-ssrA as the model substrate. Loss of GFP fluorescence is used as a reporter to monitor substrate degradation by ClpAP as a function of time (26,27). The results showed that over time, wild-type ClpP effectively degraded GFP-ssrA with a half-life of about 30 min (Fig. 4C). Meanwhile, less than a 10% decrease of the GFP-ssrA signal was observed in GFP-ssrA only group, and in wild type without ATP group, which served as negative controls. The decreasing rates of the GFP-ssrA fluorescence of ClpP_S111A_, ClpP_H136A_, and ClpP_D185A_ were significantly slower than that of ClpP_WT_, indicating the loss of their protease activity in these mutants. At the endpoint of the assay, ClpP_S111A_, ClpP_H136A_, and ClpP_D185A_ showed similar amounts of residual GFP fluorescence to the negative controls (Fig. 4C). The results suggest that the T6SS activity difference observed from ClpP_D185A_ and ClpP_S111A_, ClpP_H136A_ is not due to their protease activity difference, and the mature form of ClpP may be necessary.

### The ClpP-associated AAA+ ATPase clpA but not clpX is involved in enhancing susceptibility to A. tumefaciens T6SS activity

ClpA and ClpX are the most well-known ClpP-interacting proteins that function as the substrate recognition module of the ClpP-associated complexes. Therefore, we next determined whether Δ*clpA*::Kan (hereafter referred to as Δ*clpA*) and Δ*clpX*::Kan (hereafter referred to as Δ*clpX*) showed less susceptibility to *A. tumefaciens* T6SS-mediated antibacterial activity. Susceptibility index demonstrates that Δ*clpA* is less susceptible to *A. tumefaciens* T6SS killing than that of BW25113 wild-type (*P* = 0.02) while Δ*clpX* is similar to wild type BW25113 (*P* = 1.00) (Fig. 5A). Importantly, the decreased *A. tumefaciens* T6SS killing phenotype of Δ*clpA* can be fully complemented *in trans* (Fig. 5B). Thus, ClpA, but not ClpX, is involved in enhancing susceptibility to *A. tumefaciens* T6SS attack.

**Figure 5.**
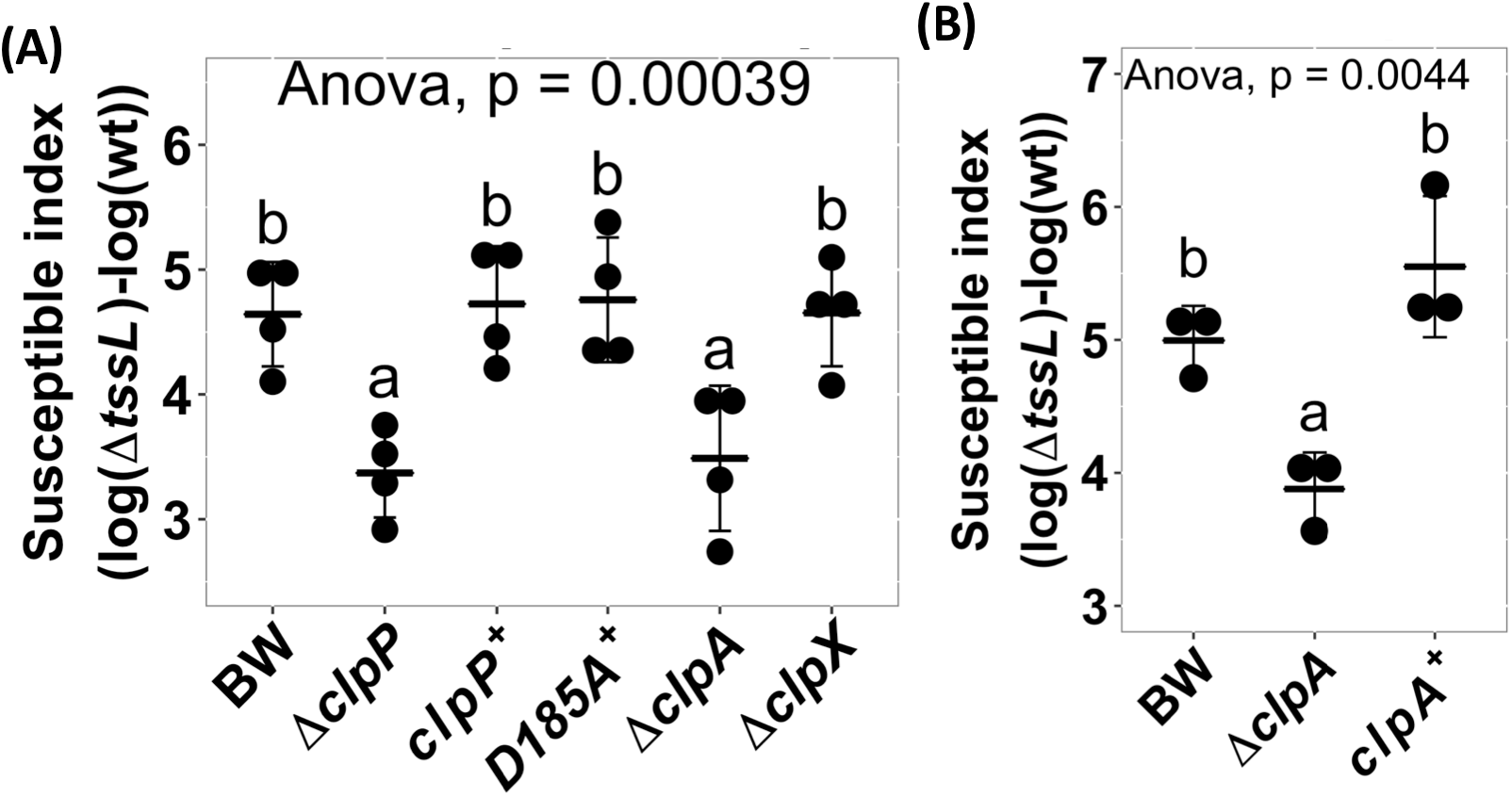
*ClpP associated AAA+ ATPase clpA but not clpX* may be involved in enhancing *A. tumefaciens* T6SS antibacterial activity. **(A)** *A. tumefaciens* T6SS antibacterial activity against *E. coli* Δ*clpP* and its complement strain, Δ*clpA* and Δ*clpX*. The *A. tumefaciens* and the *E. coli* were co-cultured at a ratio of 10:1 on AK agar medium for 16 h. Afterwards, the recovery of *E. coli* strains was quantified and the susceptible index (SI) was calculated by subtracting the difference of the recovered log(cfu) of that attacked by Δ*tssL* to that by wild type *A. tumefaciens* C58. **(B)** *A. tumefaciens* T6SS antibacterial activity assay and the SI were performed as described in (A) using *E. coli* wild type (BW), Δ*clpA*, and Δ*clpA* complemented with *clpA* expressed on plasmid (*clpA*+). Data in (A) and (B) are mean ± SD of four and three independent experiments, respectively. Statistical analysis involved single factor analysis of variance (ANOVA) and TukeyHSD with *P* < 0.05 for statistical significance.

### A ClpP variant deficient in interacting with ClpA loses its ability in enhancing E. coli susceptibility to A. tumefaciens T6SS attack

As the presence of either ClpA or ClpP resulted in enhancing *A. tumefaciens* T6SS killing, we asked whether their interaction is essential in enhancing *A. tumefaciens* T6SS killing. It has been demonstrated that the ClpP R26A and D32A variants lose their ability to bind to ClpA by 50% and 100%, respectively (25). Therefore, we complemented those mutants to Δ*clpP* to determine whether these ClpP variants could restore the susceptibility. The R26A^+^ was able to fully complement (*P* = 0.96, compared to ClpP^+^) whileD32A^+^ failed to complement and showed no statistically difference in SI than that of Δ*clpP* (*P* = 0.08) (Fig. 6). These results suggest that the interaction between ClpP and ClpA in *E. coli* is critical in enhancing the recipient cell’s susceptibility to *A. tumefaciens* T6SS antibacterial activity.

**Figure 6.**
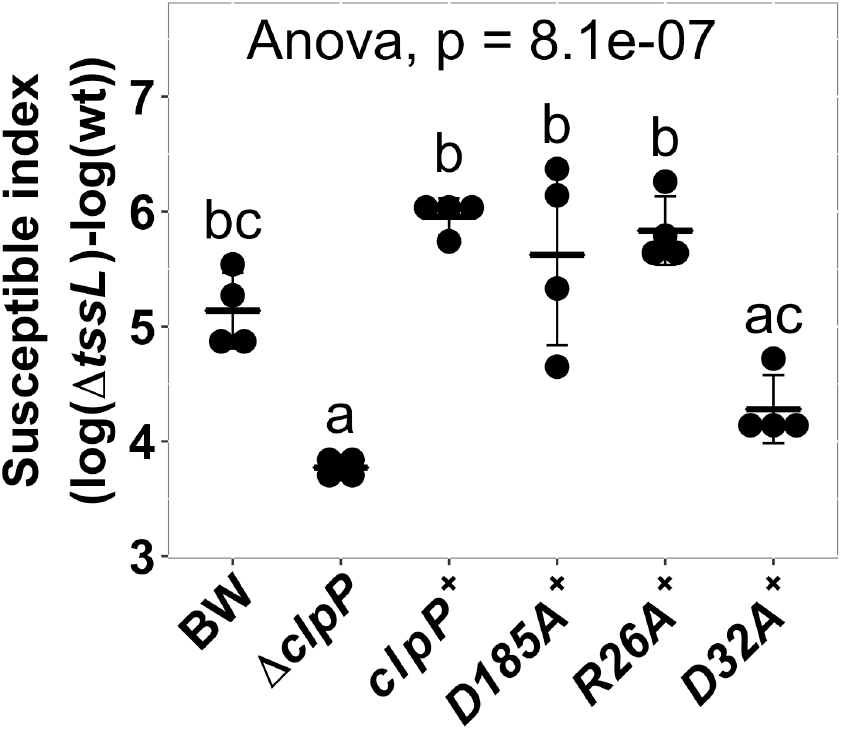
ClpP - ClpA interaction may be involved in enhancing A. tumefaciens T6SS antibacterial activity. Interbacterial competition assay between *A. tumefaciens* and *E. coli* wild type, Δ*clpP*, and Δ*clpP* complement strains expressing wild type ClpP (*clpP*^+^), catalytic mutant Clp_D185A_, ClpAP complex formation mutants ClpP_R26A_ and ClpP_D32A_. The ClpAP complex forming ability is half than that of wild type ClpP in ClpP_R26A_ and is completely loss in ClpP_D32A_ according to (25). The T6SS killing data are mean ± SD of four biological replicates from two independent experiments. Statistical analysis involved single factor analysis of variance (ANOVA) and TukeyHSD with *P* < 0.05 for statistical significance.

## Discussion

This study provided evidence that genetic factors of the recipient cells play an essential role in affecting the outcome of T6SS antibacterial activity. Using the high throughput, populational-level interbacterial competition screening platform developed in this study, we identified several recipient genetic factors other than effector-immunity pairs that can affect the outcome of *A. tumefaciens* T6SS antibacterial activity. Further exploration has led to the identification of at least six genes (*clpP*, *clpA, gltA*, *ydhS*, *ydaE*, *cbpA*) encoding known or putative cytoplasmic proteins, while CbpA is residing both in the cytoplasm and in nucleoid (28). As none of these gene products were localized to the inner membrane, periplasm, outer membrane, or extracellular region, this implies that the process affecting the outcome of *A. tumefaciens* T6SS killing to *E. coli* mainly occurs in the cytoplasm after the injection of T6SS puncturing apparatus. As exemplified by ClpP, our results further show that the universal and highly conserved ClpAP protease complex is one of the factors that enhance recipient susceptibility during *A. tumefaciens* T6SS killing. Previous studies have mainly focused on how attacker T6SS is regulated and sensed. This study provides a new insight that recipient cell genes can also affect the T6SS killing outcome and that it could take place after injection of T6SS apparatus injection into the recipient cells.

Our data showed that ClpA but not ClpX, together with ClpP, contributes to the susceptibility of the recipient *E. coli* to *A. tumefaciens* T6SS killing. The reason not having ClpX could be *clpX* transcript dropped and fade as soon as 15 minutes after the onset of carbon starvation (29), which is the condition we used for interbacterial competition. Further, ClpP variant with loss of its ability to form a complex with ClpA could not complement the phenotype as ClpP_wt_ did. This result implies that the ClpA-ClpP interacting complex, rather than ClpP alone, is the cause of the enhanced susceptibility to T6SS attack. The ClpA-ClpP complex formation starts from self-assembly of ClpA into hexameric ClpA_6_ in the presence of ATP, followed by interaction between tetradodecameric ClpP (ClpP_14_) and the ATP-bound ClpA_6_ to form a ClpAP complex (30). The ClpAP complex recognizes its substrates via ClpA, which is responsible for substrate unfolding required for subsequent protein degradation by ClpP (19,30). As the results indicated that protease activity of ClpP does not play a critical role in enhancing susceptibility to *A. tumefaciens* T6SS killing and that ClpP allosterically activates polypeptide translocation activity of ClpA (31), the necessity of ClpAP complex may depend on the ClpA unfoldase activity. The detailed mechanism on how recipient ClpAP is involved in T6SS susceptibility enhancement awaits future investigation.

The result that ClpAP complex formation but not the ClpP protease activity is the cause of the enhanced susceptibility to T6SS attack is somewhat surprising. However, hijacking a highly conserved and essential molecule of the recipient cell without hijacking their biochemical activity to improve attacker fitness is not uncommon. For example, the contact-dependent inhibition (CDI) effector CdiA^EC93^ hijacks the essential outer membrane protein BamA and inner membrane protein AcrB for its entry. BamA is one of the proteins in the BAM complex, which functions in outer membrane β-barrel proteins (OMPs) biogenesis. AcrB is an inner membrane protein that belongs to the multidrug efflux pump TolC complex. It has been demonstrated that CdiA^EC93^ hijacks BamA but not the OMPs biogenesis where it is involved, and AcrB but not the multidrug efflux function of the TolC function (32). Another CDI effector, the Cdi^EC536^, hijacks the recipient CysK and enhance the CDI-dependent killing by effector activation rather than by disrupting recipient’s cysteine synthesis ability (33). The result that the protease activity does not participate in enhancing the outcome of *A. tumefaciens* T6SS killing suggests that ClpP protease system could be a novel target that can be hijacked by attacker *A. tumefaciens* to improve its competitive advantage. It would be interesting to uncover how and what *A. tumefaciens* factor(s) hijacks this universal and highly conserved ClpP and its associated AAA^+^ ATPase substrate recognizing partner.

To our knowledge, the involvement of the ClpAP complex in enhancing recipient’s susceptibility to *A. tumefaciens* T6SS activity has not been described in the contact-dependent competitor elimination systems like T6SS, CDI, or type VII secretion system (T7SS). The ClpP protease is highly conserved in both prokaryotes and eukaryotic organelles like plastid and mitochondria. The ClpP protease cooperates with different AAA^+^ ATPases in different organisms. It works with ClpA and ClpX in gram-negative bacteria, with ClpC, and ClpE in gram-positive bacteria, and with ClpC1, ClpC2, and ClpD in the chloroplast (34,35). In all these cases, the ClpP protease seems to have a central role in protein homeostasis. Dysfunction of the system can lead to severe developmental defects, reduction in the pathogenicity, or lethal (21,36,37). The current finding is additional evidence to support that T6SS can manipulate the essential and highly conserved molecules of recipient cells to achieve better inhibiting performance (38). Elucidating the underlying molecular mechanisms of ClpAP and other recipient factors would be the next direction to further understand how genetic factors can affect the recipient susceptibility to T6SS attacks.

## Experimental procedures

### Bacterial strains, plasmids, and growth conditions

The Keio collection was used for all *E. coli* mutants, and the BW25113 strain was used as the wild type unless otherwise indicated. The complete information about the strains and plasmids used in this study are described in Table S1. *A. tumefaciens* was grown at 25 °C in 523 and *E. coli* was grown in LB at 37 °C unless indicated. The plasmids were maintained in 20 μg/mL kanamycin (Km), 100 μg/mL spectinomycin (Sp), 20 μg/mL gentamycin, for *E. coli*.

### Plasmid construction

All plasmids were confirmed by sequencing unless otherwise indicated. The complete list of primers used in this study is in Table S2. Plasmid pNptII (Table S1) was created by ligating the *Xho*I/*Bam*HI-digested *nptII* PCR product into the same restriction sites of pRL662. The plasmid was transformed into DH10B, and the resulting strain was designated as EML5395. The pRL-*rpsL*, pRL-*galK*, pRL-*nupG*, and pRL-*rpsL*^Str^ were created by ligating the *Xho*I/*Xba*I-digested PCR product into the same restriction sites of pRL662. The plasmid was transformed into DH10B, and the resulting strain was designated as EML5389, EML5390, EML5391, and EML5392, respectively. Plasmid pClpP-HA and pClpA-HA were created by ligating *Sac*I/*Pst*I-digested PCR products (*clpP* and *clpA* from BW25113 wild type without the stop codon, respectively) into pTrc200HA. The pClpP_S111A_-HA was created by amplifying fragments using pTRC99C-F plus ClpP-S111A-rv and pTRC99C-R plus ClpP-S111A-fw as primers. The two fragments were then merged and amplified by PCR-Splicing by Overlapping Extension (SOEing) (39). The resulting full-length *clpP*-containing fragment was then digested by *Sac*I/*Pst*I then ligated into pTrc200HA. All other pClpP-HA plasmids with a mutated form of ClpP were created similarly. All plasmids of pClpP-tev-His with a mutated *clpP* gene was constructed similar to that of pClpP_S111A_-HA mentioned above with the differences below: primer T7 was used instead of pTRC99C-F and primer T7T was used instead of pTRC99C-R and the restriction sites used were *Xba*I/*Xho*I.

### Whole-genome alignment

Whole-genome alignment between the chromosome of *E. coli* strain BW25113 (GenBank Accession Number CP009273) and DH10B (GenBank Accession Number CP000948) was performed using Mauve (development snapshots version 2015-02-25) (40) with default settings.

### Regular A. tumefaciens interbacterial competition assay

The optical densities of the cultured *A. tumefaciens* and *E. coli* were measured and adjusted to OD_600_ equals to 3.0 in 0.9% NaCl (w/v). The recipient *E. coli* cells were then further diluted to OD_600_ equals to 0.3 or 0.1, depending on the need of the assay. Afterward, the attacker and the recipient cultures were mixed with equal volume to make the attacker: recipient ratio 10:1 or 30:1, respectively. Ten microliters of the mixed bacterial culture were then spotted onto *<u>A</u>grobacterium* <u>K</u>ill-triggering medium (AK medium, 3 g K_2_HPO_4_, 1 g NaH_2_PO_4_, 1 g NH_4_Cl, 0.15 g KCl, 9.76 g MES, pH5.5) solidified by 2%(w/v) agar then allowed to dry to let the competition take place. The competition plates were cultured at 25°C for 16 h. After the competition, bacteria were recovered using a loop and resuspended into 500 μL 0.9% NaCl. The recovered bacterial suspension was then serially diluted and plated onto LB supplemented with spectinomycin to select recipient *E. coli* cells. After overnight culture at 37°C, the recovered colony formation unit (cfu) were counted and recorded.

### The high-throughput population level, contact-dependent antagonism screening platform

Pipetting steps of the screening platform were performed by the pipetting robot EzMate401 (Arise Biotech, Taiwan) unless otherwise specified. Fifty microliters of the cultured attacker *A. tumefaciens* was pelleted using 8,000 xg, 10 min at 15 °C. After removing the medium, the pellet was washed twice using 0.9% NaCl (w/v) then adjusted to OD_600_ equals to 3.0. The OD_600_ adjusted attacker cells were then dispensed as 300 μL into each well of a 2.2 mL-Deepwell microplate (Basic Life, Taiwan). Each well was then added by ten μL of the cultured recipient *E. coli* mutants and mixed well to make the attacker: target at 30:1 (v/v). After mixing, the bacterial mixture was then added onto the competition plate. The competition plate was made by 25 mL of the AK medium with 2%(w/v) agarose solidified in a 96-well lid. The competition plate was then cultured at 25°C for 16 h before recovery. The recovery was performed by stamping a 96-well plate replicator to the competition spots followed by resuspending the bacterial cells to a 96-well plate containing 200 μL 0.9% NaCl in each well. Ten microliters of the recovered bacterial suspension were then spotted onto LB agar supplemented with spectinomycin made in a 96-well lid, which was then cultured at 37 °C overnight before observation and image acquisition. The *E. coli* mutants that were forming multiple colonies were the candidates.

### Protein production and purification

The plasmid constructs ClpX (ClpX-ΔN-ter), wild-type ClpP and GFP-ssrA were a kind gift from Dr. Robert T. Sauer (MIT, Cambridge. The USA). Site-directed mutagenesis was performed to generate the ClpP variants. *E. coli* BL21(DE3) was used as a host to produce all proteins of interests. Cells were cultured in LB medium supplemented with appropriate antibiotics in 1 L flask. When OD_600_ reached 0.6, the bacterial culture was cooled to 16 °C, and IPTG was added (final concentration of 0.5 mM) for the over-expression of the protein. The cells were further allowed to grow for 16 h, followed by centrifugation to pellet them and resuspended in lysis buffer (50 mM Tris, pH 8.0, 300 mM Nal, 1% Triton X-100, 10 mM betamercaptoethanol, 1 mM DTT, and 10% Glycerol). The cells were lysed by sonication at 4 °C (amplitude 10 for 5 sec, followed by 15-sec breaks; total sonication time was 6 min) (Pro Scientific, USA). The lysates were centrifuged at 20,000 rpm for 30 min at 4 °C. The supernatants were collected and loaded onto Ni-NTA column (GE Healthcare, USA) equilibrated with wash buffer (50 mM Tris pH 8.0, 300 mM NaCl) and eluted by 6 mL wash buffer containing 250 mM imidazole. The eluted fractions of the protein were further subjected to size exclusion chromatography (SEC) by Superdex 200,16/60 column (GE Life science, USA) in buffer containing 50 mM Tris pH 7.5, 100 mM KCl, 25 mM MgCl_2_ 1 mM DTT, 10% glycerol. The protein purity was confirmed on 12% SDS-PAGE. The samples were flash-frozen and stored in −80 °C until further use.

### Protein degradation assay

GFP fluorescence based-degradation assays were carried out in PD buffer (25 mM HEPES, pH 7.5, 100 mM KCl, 25 mM MgCl_2_, 1 mM DTT, 10% glycerol) containing 3 μM GFP-ssrA as substrate and ATP regeneration system (16 mM creatine phosphatase, 0.32 mg/mL creatine kinase) as described previously (27). In brief, 0.1 μM ClpX_6_ and 0.3 μM ClpP_14_ or its variants were mixed at 30 °C and let stand for 2 min. The protein degradation reaction was started by addition of ATP to a final concentration of 5 mM. The changes in the fluorescence were measured at 511 nm with an excitation wavelength at 467 nm in a 96-well format using Infinite M1000 PRO plate reader (TECAN, Switzerland).

### Statistical analysis

Statistical analyses were performed using the R program (version 3.5.1). One-way analysis of variance (one-way ANOVA) and Tukey’s honestly significant difference test (Tukey HSD), in which significant difference threshold set as 0.05, were used in all case.

## Acknowledgments

The authors thank National BioResource Project (NIG, Japan): *E.coli* for providing the Keio collection. We acknowledge the staffs in the Plant Cell Biology Core Laboratory and the DNA Sequencing Core Laboratory at the Institute of Plant and Microbial Biology, and Biophysics Facility at the Institute of Biological Chemistry, Academia Sinica, Taiwan, for technical support.

## Conflict of interests

The authors declare that they have no conflicts of interest with the content of this article.

## FOOTNOTES

Funding for this project was provided by the Ministry of Science and Technology of Taiwan (MOST) (grant no. 104-2311-B-001-025-MY3 to E.-M.L; 107-2628-M-001-005-MY3 to S.-T.D.H; 107-2811-M-001-1574 to M.K.S.) and Academia Sinica Investigator Award to E.-M.L. (grant no. AS-IA-107-L01).

## abbreviations

T6SS: type VI secretion system;
SI: susceptible index;
E-I: effector-immunity;
CDI: contact-dependent inhibition;
T7SS: type VII secretion system

## References

1. Stubbendieck, R. M., Vargas-Bautista, C., and Straight, P. D. (2016) Bacterial communities: Interactions to scale. Front. Microbial. 7 10.3389/fmicb.2016.01234

2. Hacquard, S., Garrido-Oter, R., González, A., Spaepen, S., Ackermann, G., Lebeis, S., McHardy, Alice C., Dangl, Jeffrey L., Knight, R., Ley, R., and Schulze-Lefert, P. (2015) Microbiota and host nutrition across plant and animal kingdoms. Cell Host & Microbe 17, 603–616

3. Foster, Kevin R., and Bell, T. (2012) Competition, not cooperation, dominates interactions among culturable microbial species. Curr. Biol. 22, 1845–1850

4. Ghoul, M., and Mitri, S. (2016) The ecology and evolution of microbial competition. Trends Microbiol. 24, 833–845

5. Coulthurst, S. J. (2013) The type VI secretion system – a widespread and versatile cell targeting system. Res. Microbiol. 164, 640–654

6. Lien, Y.-W., and Lai, E.-M. (2017) Type VI secretion effectors: Methodologies and biology. Front. in Cell. Infect. Microbiol. 7 10.3389/fcimb.2017.00254

7. Ho, Brian T., Dong, Tao G., and Mekalanos, John J. (2014) A view to a kill: The bacterial type VI secretion system. Cell Host & Microbe 15, 9–21

8. Bernal, P., Allsopp, L. P., Filloux, A., and Llamas, M. A. (2017) The *Pseudomonas putida* T6SS is a plant warden against phytopathogens. ISME J. 11, 972–987

9. Sana, T. G., Lugo, K. A., and Monack, D. M. (2017) T6SS: The bacterial “fight club” in the host gut. PLoS Path. 13, e1006325

10. Basler, M., Ho, B. T., and Mekalanos, J. J. (2013) Tit-for-tat: Type VI secretion system counterattack during bacterial cell-cell interactions. Cell 152, 884–894

11. Dong, T. G., Ho, B. T., Yoder-Himes, D. R., and Mekalanos, J. J. (2013) Identification of T6SS-dependent effector and immunity proteins by Tn-seq in *Vibrio cholerae*. PNAS 110, 2623–2628

12. Ma, L.-S., Hachani, A., Lin, J.-S., Filloux, A., and Lai, E.-M. (2014) *Agrobacterium tumefaciens* deploys a superfamily of type VI secretion dnase effectors as weapons for interbacterial competition *in planta*. Cell Host & Microbe 16, 94–104

13. Sana, T. G., Flaugnatti, N., Lugo, K. A., Lam, L. H., Jacobson, A., Baylot, V., Durand, E., Journet, L., Cascales, E., and Monack, D. M. (2016) *Salmonella* typhimurium utilizes a T6SS-mediated antibacterial weapon to establish in the host gut. PNAS 113, E5044–E5051

14. Wu, C.-F., Lin, J.-S., Shaw, G.-C., and Lai, E.-M. (2012) Acid-induced type VI secretion system is regulated by ExoR-ChvG/ChvI signaling cascade in *Agrobacterium tumefaciens*. PLoS Pathog. 8, e1002938

15. Wu, C.-F., Santos, M. N. M., Cho, S.-T., Chang, H.-H., Tsai, Y.-M., Smith, D. A., Kuo, C.-H., Chang, J., and Lai, E.-M. (2019) Plant pathogenic *Agrobacterium tumefaciens* strains have diverse type VI effector-immunity pairs and vary in *in planta* competitiveness. Mol. Plant-Microbe Interact. 10.1094/MPMI-01-19-0021-R

16. Whitney, John C., Quentin, D., Sawai, S., LeRoux, M., Harding, Brittany N., Ledvina, Hannah E., Tran, Bao Q., Robinson, H., Goo, Young A., Goodlett, David R., Raunser, S., and Mougous, Joseph D. (2015) An interbacterial NAD(P) ^+^ glycohydrolase toxin requires Elongation Factor Tu for delivery to target cells. Cell 163, 607–619

17. Quentin, D., Ahmad, S., Shanthamoorthy, P., Mougous, J. D., Whitney, J. C., and Raunser, S. (2018) Mechanism of loading and translocation of type VI secretion system effector Tse6. Nat. Microbiol. 3, 1142–1152

18. Mariano, G., Monlezun, L., and Coulthurst, S. J. (2018) Dual role for DsbA in attacking and targeted bacterial cells during type VI secretion system-mediated competition. Cell Rep. 22, 774–785

19. Olivares, A. O., Baker, T. A., and Sauer, R. T. (2015) Mechanistic insights into bacterial AAA+ proteases and protein-remodelling machines. Nat. Rev. Microbiol. 14, 33

20. Mahmoud, S. A., and Chien, P. (2018) Regulated proteolysis in bacteria. Annu. Rev. Biochem. 87, 677–696

21. Bhandari, V., Wong, K. S., Zhou, J. L., Mabanglo, M. F., Batey, R. A., and Houry, W. A. (2018) The role of ClpP protease in bacterial pathogenesis and human diseases. ACS Chem. Biol. 13, 1413–1425

22. Alexopoulos, J. A., Guarné, A., and Ortega, J. (2012) Clpp: A structurally dynamic protease regulated by AAA+ proteins. J. Struct. Biol. 179, 202–210

23. Wang, J., Hartling, J. A., and Flanagan, J. M. (1997) The structure of ClpA at 2.3 Å resolution suggests a model for ATP-dependent proteolysis. Cell 91, 447–456

24. Maurizi, M. R., Clark, W. P., Kim, S. H., and Gottesman, S. (1990) Clp P represents a unique family of serine proteases. J. Biol. Chem. 265, 12546–12552

25. Bewley, M. C., Graziano, V., Griffin, K., and Flanagan, J. M. (2006) The asymmetry in the mature amino-terminus of ClpP facilitates a local symmetry match in ClpAP and ClpXP complexes. J. Struct. Biol. 153, 113–128

26. Weber-Ban, E. U., Reid, B. G., Miranker, A. D., and Horwich, A. L. (1999) Global unfolding of a substrate protein by the Hsp100 chaperone ClpA. Nature 401, 90–93

27. Sriramoju, M. K., Chen, Y., Lee, Y.-T. C., and Hsu, S.-T. D. (2018) Topologically knotted deubiquitinases exhibit unprecedented mechanostability to withstand the proteolysis by an AAA+ protease. Sci. Rep. 8, 7076

28. Orfanoudaki, G., and Economou, A. (2014) Proteome-wide subcellular topologies of *E. coli* polypeptides database (STEPdb). Mol. Cell. Proteom.: MCP 13, 3674–3687

29. Li, C., Tao, Y. P., and Simon, L. D. (2000) Expression of different-size transcripts from the ClpP-ClpX operon of *Escherichia coli* during carbon deprivation. J. Bacteriol. 182, 6630–6637

30. Maurizi, M. R. (1991) ATP-promoted interaction between Clp A and Clp P in activation of Clp protease from *Escherichia coli*. Biochem. Soc. Trans. 19, 719–723

31. Miller, J. M., Lin, J., Li, T., and Lucius, A. L. (2013) *E. coli* ClpA catalyzed polypeptide translocation is allosterically controlled by the protease ClpP. J. Mol. Biol. 425, 2795–2812

32. Aoki, A. S. K., C., M. J., Kyle, J., Benjamin, T., Rupinderjit, P., N., T. B., Susan, R., Julia, W., A., B. B., J., S. T., and A., L. D. (2008) Contact-dependent growth inhibition requires the essential outer membrane protein BamA (YeaT) as the receptor and the inner membrane transport protein acrb. Mol. Microbiol. 70, 323–340

33. Diner, E. J., Beck, C. M., Webb, J. S., Low, D. A., and Hayes, C. S. (2012) Identification of a target cell permissive factor required for contact-dependent growth inhibition (CDI). Genes Dev. 26, 515–525

34. Moreno, J. C., Tiller, N., Diez, M., Karcher, D., Tillich, M., Schöttler, M. A., and Bock, R. (2017) Generation and characterization of a collection of knock-down lines for the chloroplast Clp protease complex in tobacco. J. Exp. Bot. 68, 2199–2218

35. Culp, E., and Wright, G. D. (2016) Bacterial proteases, untapped antimicrobial drug targets. J. Antibiot. 70, 366–377

36. Nishimura, K., and van Wijk, K. J. (2015) Organization, function and substrates of the essential Clp protease system in plastids. Biochim. Biophys. Acta 1847, 915–930

37. Cole, A., Wang, Z., Coyaud, E., Voisin, V., Gronda, M., Jitkova, Y., Mattson, R., Hurren, R., Babovic, S., Maclean, N., Restall, I., Wang, X., Jeyaraju, Danny V., Sukhai, Mahadeo A., Prabha, S., Bashir, S., Ramakrishnan, A., Leung, E., Qia, Yi H., Zhang, N., Combes, Kevin R., Ketela, T., Lin, F., Houry, Walid A., Aman, A., Al-awar, R., Zheng, W., Wienholds, E., Xu, Chang J., Dick, J., Wang, Jean C. Y., Moffat, J., Minden, Mark D., Eaves, Connie J., Bader, Gary D., Hao, Z., Kornblau, Steven M., Raught, B., and Schimmer, Aaron D. (2015) Inhibition of the mitochondrial protease ClpP as a therapeutic strategy for human acute myeloid leukemia. Cancer Cell 27, 864–876

38. Russell, A. B., Peterson, S. B., and Mougous, J. D. (2014) Type VI secretion system effectors: Poisons with a purpose. Nat Rev Micro 12, 137–148

39. Heckman, K. L., and Pease, L. R. (2007) Gene splicing and mutagenesis by PCR-driven overlap extension. Nat. Protoc. 2, 924–932

40. Darling, A. C. E., Mau, B., Blattner, F. R., and Perna, N. T. (2004) Mauve: Multiple alignment of conserved genomic sequence with rearrangements. Genome Res. 14, 1394–1403

41. Keseler, I. M., Mackie, A., Santos-Zavaleta, A., Billington, R., Bonavides-Martínez, C., Caspi, R., Fulcher, C., Gama-Castro, S., Kothari, A., Krummenacker, M., Latendresse, M., Muñiz-Rascado, L., Ong, Q., Paley, S., Peralta-Gil, M., Subhraveti, P., Velázquez-Ramírez, D. A., Weaver, D., Collado-Vides, J., Paulsen, I., and Karp, P. D. (2016) The EcoCyc database: Reflecting new knowledge about *Escherichia coli* K-12. Nucleic Acids Res. 45, D543–D550

