## Supplemental Table 1, Supplemental Table 2, Supplemental Figure 1 for "Identification of *Escherichia coli* ClpAP in regulating susceptibility to type VI secretion system-mediated attack by *Agrobacterium tumefaciens*"

**Supporting Information**

Table S1. Bacterial strains and plasmids
Table S2. Primer information
Fig. S1

Table S1. Bacterial strains and plasmids

| Strain/Plasmid | Relevant characteristics | Source/reference |
|---|---|---|
| *A. tumefaciens* | | |
| C58 (EML530) | wild type virulence strain containing pTiC58 and pAtC58 | Eugene Nester |
| C58::Δ*atu4333* (EML1073) | *atu4333* (*tssL*) in-frame deletion mutant of C58 background | (1) |
| *E. coli* | | |
| DH10B | Host for DNA cloning | Invitrogen |
| BW25113 | wild type strain of the Keio Collection. *rrnB* DE*lacZ*4787 *HsdR*514 DE(*araBAD*)567 DE(*rhaBAD*)568 *rph*-1. | (2) |
| Keio collection | Systematic single-gene knock-out mutants of *E. coli* BW25113 | (2) |
| JW0427 | BW25113 *clpP*::*kan* | (2) |
| JW0866 | BW25113 *clpA*::*kan* | (2) |
| JW0428 | BW25113 *clpX*::*kan* | (2) |
| EML5395 | DH10B harboring pNptII | This study |
| EML5393 | BW25113 wild type harboring pNptII | This study |
| BL21(DE3) | Host for protein expression | (3) |
| Plasmids | | |
| pTrc200HA | $Sp^R$, pTrc200 harboring C-terminal influenza hemagglutinin (HA) epitope, P*trc*, $lacI^q$, pVS1 origin | Laboratory collection |
| pRL662 | $Gm^R$, a non-transferable broad-host range vector derived from pBBR1MCS2 | (4) |

| | | |
|---|---|---|
| pET22b(+) | $Ap^R$, *E. coli* overexpression vector harboring C-terminal 6xHis epitope | Novagen |
| pRL-*rpsL* | $Gm^R$, pRL662 expressing BW25113 *rpsL* gene | This study |
| pRL-*galk* | $Gm^R$, pRL662 expressing BW25113 *galK* gene | This study |
| pRL-*nupG* | $Gm^R$, pRL662 expressing BW25113 *nupG* gene | This study |
| pRL-*rpsL*$^{Str}$ | $Gm^R$, pRL662 expressing DH10B *rpsL*$^{Str}$ gene | This study |
| pNptII | $Km^R$, $Gm^R$, pRL662 expressing *nptII* gene | This study |
| pClpP-HA | $Sp^R$, pTrc200HA expressing ClpP-HA fusion protein | This study |
| pClpA-HA | $Sp^R$, pTrc200HA expressing ClpA-HA fusion protein | This study |
| pClpP$_{S111A}$-HA | $Sp^R$, pTrc200HA expressing ClpP-HA fusion protein with S111A substitution | This study |
| pClpP$_{H136A}$-HA | $Sp^R$, pTrc200HA expressing ClpP-HA fusion protein with H136A substitution | This study |
| pClpP$_{D185A}$-HA | $Sp^R$, pTrc200HA expressing ClpP-HA fusion protein with D185A substitution | This study |
| pClpP$_{R26A}$-HA | $Sp^R$, pTrc200HA expressing ClpP-HA fusion protein with R26A substitution | This study |
| pClpP$_{D32A}$-HA | $Sp^R$, pTrc200HA expressing ClpP-HA fusion protein with D32A substitution | This study |
| pClpP-tev-His | $Ap^R$, pET22b(+) expressing ClpP-tev-His, in which ClpP protein is fused with a TEV protease cleavage site and a His-tag in its C-terminal | This study |
| pClpP$_{S111A}$-tev-His | $Ap^R$, pET22b(+) expressing ClpP-tev-His with ClpP S111A substitution | This study |
| pClpP$_{H136A}$-tev-His | $Ap^R$, pET22b(+) expressing ClpP-tev-His with ClpP H136A substitution | (5) |
| pClpP$_{D185A}$-tev-His | $Ap^R$, pET22b(+) expressing ClpP-tev-His with ClpP D185A substitution | This study |

Table S2. Primer information

| Primer | Sequence (5'-3')[a] | Plasmids |
|---|---|---|
| T7 | TAA TAC GAC TCA CTA TAG GG | pET22b(+) |
| T7T | GCT AGT TAT TGC TCA GCG G | |
| pTRC99C-F | TTGCGCCGACATCATAAC | pTrc200HA |
| pTRC99C-R | CTGCGTTCTGATTTAATCTG | |
| rpsL-fw | AAAAACTCGAGGCAAAAGCTAAAACCAGGA | 1. pRL-*rpsL* |
| rpsL-rv | AAAAATCTAGACTTACTTAACGGAGAACCA | 2. pRL-rpsL$^{Str}$ |
| galK-fw | AAAAACTCGAGCAGTCAGCGATATCCATT | pRL-galk |
| galK-rv | AAAAATCTAGAGCAAAGTTAACAGTCGGT | |
| nupG-fw | AAAAACTCGAGTCAAACACTCATCCGCAT | pRL-nupG |
| nupG-rv | AAAAATCTAGACCCGTTTTTCTTTGCGTAA | |
| NptII-fw-XhoI | AAAAACTCGAGAGACTGGGCGGTTTTATGGA | pNptII |
| NptII-rv-HindIII | AAAAAAAGCTTCTCTAGCGAACCCCAGAGTC | |
| ClpP-SacI-fw | AAAAAGAGCTCATGTCATACAGCGGCGAACGAGATAAC | pClpP-HA |
| ClpP-PstI-rv | AAAAACTGCAGATTACGATGGGTCAGAATCGAATCGAC | |
| ClpA-SacI-fw | AAAAAGAGCTCATGCTCAATCAAGAACTGGAACTCAGTTT | pClpA-HA |
| ClpA-PstI-rv | AAAAACTGCAGATGCGCTGCTTCCGCCTTGTGCTTT | |
| ClpP-S111A-fw | TGTATGGGCCAGGCGGCC**GCG**ATGGGCGCTTTCTTGCTG | 1. pClpP$_{S111A}$-HA |
| ClpP-S111A-rv | CAGCAAGAAAGCGCCCAT**CGC**GGCCGCCTGGCCCATACA | 2. pClpP$_{S111A}$-tev-His |
| ClpP-H136A-fw | AATTCGCGCGTGATGATT**GCC**CAACCGTTGGGCGGCTAC | 1. pClpP$_{H136A}$-HA |
| ClpP-H136A-rv | GTAGCCGCCCAACGGTTG**GGC**AATCATCACGCGCGAATT | 2. pClpP$_{H136A}$-tev-His |
| ClpP-D185A-fw | GAACGTGATACCGAGCGC**GCT**CGCTTCCTTTCCGCCCCT | 1. pClpP$_{D185A}$-HA |
| ClpP-D185A-rv | AGGGGCGGAAAGGAAGCG**AGC**GCGCTCGGTATCACGTTC | 2. pClpP$_{D185A}$-tev-His |
| ClpP-R26A-fw | GTCATTGAACAGACCTCA**GCC**GGTGAGCGCTCTTTTGAT | pClpP$_{R26A}$-HA |
| ClpP-R26A-rv | ATCAAAAGAGCGCTCACC**GGC**TGAGGTCTGTTCAATGAC | |
| ClpP-D32A-fw | CGCGGTGAGCGCTCTTTT**GCT**ATCTATTCTCGTCTACTT | pClpP$_{D32A}$-HA |
| ClpP-D32A-rv | AAGTAGACGAGAATAGAT**AGC**AAAAGAGCGCTCACCGCG | |

a: Restriction enzyme sites are underlined, and mutated sequences are indicated by bold type.

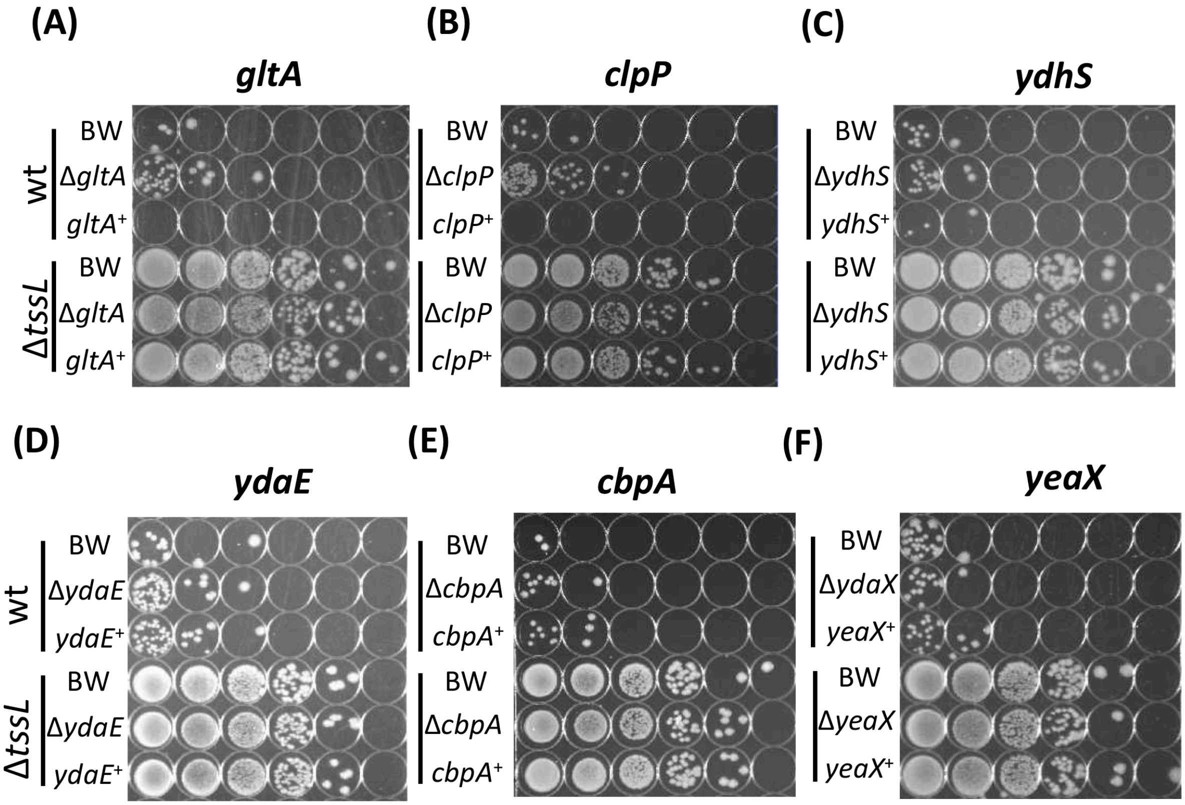


**Fig. S1 Regular interbacterial competition assay between *A. tumefaciens* C58 and the *E. coli* candidates that were less susceptible to T6SS killing.** The *A. tumefaciens* C58 wild-type (wt) or Δ*tssL* were co-cultured at a ratio of 30:1 with *E. coli* BW25113 wild type (BW), the Keio mutant strains, and the Keio mutant strain complemented with respective disrupted gene expressed on plasmid. The Keio strain used was **(A)** Δ*gltA*, **(B)** Δ*clpP*, **(C)** Δ*ydhS*, **(D)** Δ*ydaE*, **(E)** Δ*cbpA*, and **(F)** Δ*yeaX*. Data shown are representative results, at least two independent experiments were performed in each group.

References

1. Ma, L.-S., Narberhaus, F., and Lai, E.-M. (2012) IcmF family protein TssM exhibits ATPase activity and energizes type VI secretion. *J. Biol. Chem.* **287**, 15610-15621
2. Baba, T., Ara, T., Hasegawa, M., Takai, Y., Okumura, Y., Baba, M., Datsenko, K. A., Tomita, M., Wanner, B. L., and Mori, H. (2006) Construction of *Escherichia coli* K-12 in-frame, single-gene knockout mutants: The Keio collection. *Mol. Syst. Biol.* **2**, 2006.0008-10.1038/msb4100050
3. Studier, F. W., and Moffatt, B. A. (1986) Use of bacteriophage T7 RNA polymerase to direct selective high-level expression of cloned genes. *J. Mol. Biol.* **189**, 113-130
4. Vergunst, A. C., Schrammeijer, B., den Dulk-Ras, A., de Vlaam, C. M. T., Regensburg-Tuïnk, T. J. G., and Hooykaas, P. J. J. (2000) VirB/D4-dependent protein translocation from *Agrobacterium* into plant cells. *Science* **290**, 979-982
5. Sriramoju, M. K., Chen, Y., Lee, Y.-T. C., and Hsu, S.-T. D. (2018) Topologically knotted deubiquitinases exhibit unprecedented mechanostability to withstand the proteolysis by an AAA+ protease. *Sci. Rep.* **8**, 7076 10.1038/s41598-018-25470-0
